# Metagenomic analysis suggests broad metabolic potential in extracellular symbionts of the bivalve *Thyasira* cf. *gouldi*

**DOI:** 10.1101/330373

**Authors:** Bonita McCuaig, Lourdes Peña-Castillo, Suzanne C. Dufour

## Abstract

Next-generation sequencing has opened new avenues for studying metabolic capabilities of bacteria that cannot be cultured. Here, we provide a metagenomic description of a chemoautotrophic gammaproteobacterial symbiont associated with *Thyasira* cf. *gouldi*, a sediment-dwelling bivalve from the family Thyasiridae. Symbionts of thyasirids differ from those of other bivalves by being located outside rather than inside gill epithelial cells, and recent work suggests that they are capable of living freely in the environment. The *T*. cf. *gouldi* symbiont genome shows no signs of genomic reduction and contains many genes that would only be useful outside the host, including flagellar and chemotaxis genes. The thyasirid symbiont may be capable of sulfur oxidation via both the sulfur oxidation and dissimilatory sulfate reduction pathways, as observed in other bivalve symbionts. In addition, genes for hydrogen oxidation and dissimilatory nitrate reduction were found, suggesting varied metabolic capabilities under a range of redox conditions. The genes of the tricarboxylic acid cycle are also present, along with membrane bound sugar importer channels, suggesting that the bacteria may be mixotrophic. In this study, we have generated the first thyasirid symbiont genomic resources and lay the groundwork for further research in tracking the changes required for life as a bivalve symbiont.

## Introduction

Many species of marine bivalves living near oxic-anoxic boundaries form nutritional symbioses with chemoautotrophic bacteria, which are maintained in or on the host’s gills (1-4). In such associations, called chemosynthetic symbioses or chemosymbioses, the bacteria provide the host with nutrients and protection from chemical stress, while the host constitutes a protective and suitable environment for the bacterial symbionts (2, 5, 6). The metabolism of symbionts allows hosts to colonize new and often nutrient-poor niches and contributes to their ecological and evolutionary success, as moving into a niche with less competition for resources can lead to evolutionary radiation (7, 8).

Symbionts can be acquired by new generations of hosts in various ways. Vertical transmission is the transfer of bacteria from one generation to the next through gametes, most commonly the eggs; through this pathway, symbionts tend to become obligate. In horizontal transmission, host larvae are inoculated by symbionts released by nearby adults, whereas in environmental transmission, juveniles are inoculated from a free-living symbiont population (6). While the latter mode of transmission does not guarantee symbiont transfer, it does confer some advantages to both partners. Bacteria can avoid genomic reduction (i.e. the deletion of genes that no longer improve symbiont fitness) by maintaining a free-living population. Once genomic reduction occurs (via vertical transmission), the bacteria cannot survive without the host, and become reliant upon them (9); this does not occur in symbionts that maintain a functional environmental population. The maintenance of variation within bacterial populations can also benefit the host, which can be inoculated by symbionts that are well adapted for the local environment, and not necessarily the strain that their parents hosted. As the relationship between symbiotic partners becomes tighter, it may become obligate for both parties. In the case of the host, obligate symbioses can result in reduction or loss of the digestive tract, as nutritional reliance upon symbionts increases (6).

The symbiont can supplement nutrients that are lacking in the host’s diet, or simply provide an additional source of nutrients. The mode of nutrient transfer from symbiont to host varies by relationship, and in many cases, is not well defined. Some symbionts have been shown to actively transfer nutrients to their host, others have “leaky membranes” that allow nutrients to escape the bacteria, and in other cases the host consumes the bacteria through phagocytosis (10, 11). Some metabolic cycles of the symbionts may remove toxins present in the environment, providing the host protection from these compounds (12, 13). The sox and dsr cycles may remove toxic sulfur compounds while providing energy for carbon fixation. The nitrite reduction (nir) pathway removes harmful nitrogen compounds by using them as an electron sink, but this process is not always coupled with carbon fixation (14). One approach to examining the metabolic potential of chemoautotrophic symbionts is to perform genomic, or metagenomic sequencing (15-18). By identifying key genes in sequencing data, we can make inferences about the metabolic capabilities of the symbiont.

The bivalve genus *Thyasira* (Family Thyasiridae) contains both symbiotic and asymbiotic species, a seemingly unique condition among bivalve genera (19, 20). In contrast to other clams, thyasirids maintain their symbionts among the microvilli of gill epithelial cells, as described in some mussels; such extracellular symbioses have been considered more primitive than intracellular symbioses (6, 20-23). Chemosymbiotic thyasirids are mixotrophs that appear to rely on particulate food to a greater extent when symbiont abundance is low (24), or at times when environmental sulfide concentrations are low (25). All thyasirid symbionts identified to date are gammaproteobacteria (23, 25, 26). The thyasirid symbionts are clustered into divergent groups which include both symbiotic and free-living sulfur-oxidizing bacteria (23). Enzymatic and PCR techniques have shown the presence of ribulosebisphosphate carboxylase (RuBisCO) and adenylylsulphate reductase in the symbionts of all chemosymbiotic thyasirids investigated (23, 25).

In Bonne Bay, Newfoundland, Canada, gammaproteobacteria have been found living extracellularly on the gills of thyasirid clams identified as *Thyasira* cf. *gouldi* OTUs 1 and 2 (20). Phylogenetic analysis using 16S rRNA sequences have identified three distinct symbiont phylotypes (A – C) hosted by the two *T.* cf. *gouldi* OTUs (27, 28). There was no apparent co-speciation between host and symbiont as both clam OTUs could host any one of the three symbiont phylotypes, and there is some evidence of multiple strains infecting a single host, although this has not been confirmed (27, 28). The three bacterial 16S rRNA phylotypes cluster during phylogenic analysis and are closely related to the *Thyasira flexuosa* symbiont and to tubeworm symbionts (notably those associated with *Riftia pachyptila*) as well as free-living sulfur oxidizing bacteria (27, 28). *T.* cf. *gouldi* symbionts have been identified within surrounding sediment samples, supporting an environmental mode of transmission and the existence of a free-living symbiont population (29).

We present here the first genomic analysis of a thyasirid symbiont, that of *T.* cf. *gouldi* symbiont phylotype B (one of the most common; 27, 28). This investigation is of particular interest given the extracellular location and facultative nature of thyasirid symbionts, and provides a contrast to genomic studies of intracellular (and often obligate) bivalve chemosymbionts. After providing an overview of the metagenomic data collected, we characterize important metabolic cycles, including carbon fixation and sulfur oxidation, and identify genomic characteristics that allow us to infer the mode of symbiont transmission and support the evidence for a free-living state in thyasirid bacterial symbionts. By identifying the genes for metabolic pathways in symbiont genomes, we lay the groundwork for future transcription and protein studies.

## Results and Discussion

### Genomic overview

Sequence reads can be found on the SRA database under sample SRS1569030, sequencing runs SRR3928943 and SRR3928944. Assembled contigs were uploaded to GenBank under BioProject PRJNA327811, accession number SAMN05358035. The assembly resulted in a highly fractured draft genome, suggesting heterogeneity in the bacterial population (recently suggested in reference 28) as genes assembled well, but intergenic spaces did not. Nevertheless, the symbiont population in the *T.* cf. *gouldi* specimen studied was comprised of a single species, as only one complete 16S rRNA sequence was found. A similarly fragmented genome despite the presence of a single 16S rRNA sequence was described in a metagenomic study of *R. pachyptila* symbionts, which are environmentally acquired (16).

The assembly resulted in 12,504 contigs, with an N50 of 1500. The GC content is 42 ± 7%. In total, 20,843 putative genes were assembled and possible functions were assigned to 3,339 of them allowing us to infer some of the metabolic capabilities of the *T.* cf. *gouldi* symbiont. A summary of Level 1 Subsystem Functions is presented in Table 1.

**Table 1.** The number of putative proteins assigned to level 1 subsystem functions

| Level 1 Subsystem Functions | Number of genes assigned |
| --- | --- |
| Amino Acids and Derivatives | 211 |
| Carbohydrates | 177 |
| Miscellaneous | 151 |
| DNA Metabolism | 136 |
| Protein Metabolism | 148 |
| Cell Wall and Capsule | 113 |
| Cofactors, Vitamins, Prosthetic Groups, Pigments | 109 |
| RNA Metabolism | 107 |
| Regulation and Cell signaling | 106 |
| Respiration | 102 |
| Membrane Transport | 90 |
| Stress Response | 66 |
| Virulence, Disease and Defense | 61 |
| Nitrogen Metabolism | 54 |
| Phages, Prophages, Transposable elements, Plasmids | 42 |
| Fatty Acids, Lipids, and Isoprenoids | 40 |
| Motility and Chemotaxis | 38 |
| Nucleosides and Nucleotides | 38 |
| Sulfur Metabolism | 30 |
| Phosphorus Metabolism | 25 |
| Cell Division and Cell Cycle | 25 |
| Metabolism of Aromatic Compounds | 22 |
| Other | 382 |

### Genomic support of environmental transmission

Some general characteristics of the genome support the capability of *T.* cf. *gouldi* symbionts to have a free-living existence: there is no sign of genomic reduction (as discussed below), the GC content (48%) is similar to that of environmental bacteria (those that have undergone genome reduction often have GC contents below 40%), and mobile elements are present (30). While none of these are conclusive evidence of environmental transmission, they are uncommon in co-evolved vertically transmitted symbionts (30); other evidence, such as the identification of the symbiont 16S rRNA gene in surrounding sediments (29), supports a free-living capability.

Four different mobile elements were identified using the MG-RAst website and PROKKA (see Methods), although the exact number of copies was unclear because of the fragmented nature of the draft genome. Phage genes were also identified, although the exact number was again indiscernible. There was no apparent loss of genes for DNA repair in the *T.* cf. *gouldi* symbiont genome (Table 2), in contrast to the reduced genome of the vesicomyid symbionts which lacks *rec*A for genetic recombination and *mut*Y for DNA repair (31). Genes for DNA repair are often lost in vertically transmitted symbionts, contributing to GC bias and the presence of many pseudogenes (30, 31).

**Table 2:** Putative gene functions associated with the Calvin-Benson-Bassham Cycle

| Putative gene function | Number of copies in draft genome | Percent identity | Alignment length (bp) |
| --- | --- | --- | --- |
| Fructose-bisphosphate aldolase class I | 1 | 78.93 | 337 |
| Fructose-bisphosphate aldolase class II | 1 | 75.42 | 354 |
| NAD-dependent glyceraldehyde-3-phosphate dehydrogenase | 2 | 80.39 | 225.5 |
| NADPH-dependent glyceraldehyde-3-phosphate dehydrogenase | 1 | 83.54 | 328 |
| Phosphoglycerate kinase | 1 | 71.58 | 285 |
| Phosphoribulokinase | 2 | 80.21 | 96 |
| Ribose 5-phosphate isomerase A | 4 | 73.16 | 71.75 |
| Ribulose bisphosphate carboxylase | 2 | 83.3 | 180 |
| Ribulose-phosphate 3-epimerase | 1 | 84.68 | 222 |
| Transketolase | 1 | 71.61 | 384 |
| Transketolase, C-terminal section | 2 | 62.81 | 112.5 |
| Transketolase, N-terminal section | 1 | 61.73 | 81 |
| Triosephosphate isomerase | 1 | 67.07 | 167 |

One contig contained many plasmid related genes, and further analysis uncovered the presence of a circular extrachromosomal plasmid. A *vir*B operon consisting of 10 genes encoding a type IV secretion system was identified on the putative plasmid (Figure 1). The type IV secretion system can be used in conjunction with pili for conjugation, however, the virB operon can also be critical in both pathogenic and mutualistic relationships as a secretion system that moves molecules from bacteria to the host (32). In many mutualistic relationships, these molecules act to identify, colonize, and communicate with the host in a non-harmful way, with the most common molecules moved across cell walls by this secretion system being DNA (33, 34). However, in different bacteria the system can transport a number of different molecules; notably, in some pathogenic species it can transfer small effector proteins (33). The genes *vir*B1-5 are often transcribed together and *vir*B7-11 form another co-transcribed group (34). On the *T.* cf. *gouldi* symbiont plasmid, the *vir*B genes show a similar arrangement, with two hypothetical proteins placed between *vir*B5 and *vir*B6 (Fig. 1). Effector sequences were not identified, and no putative function was found for the nine hypothetical proteins on the plasmid. The importance of this plasmid is unknown, and the secretion system may not be important in the symbiosis, but simply used for conjugation.

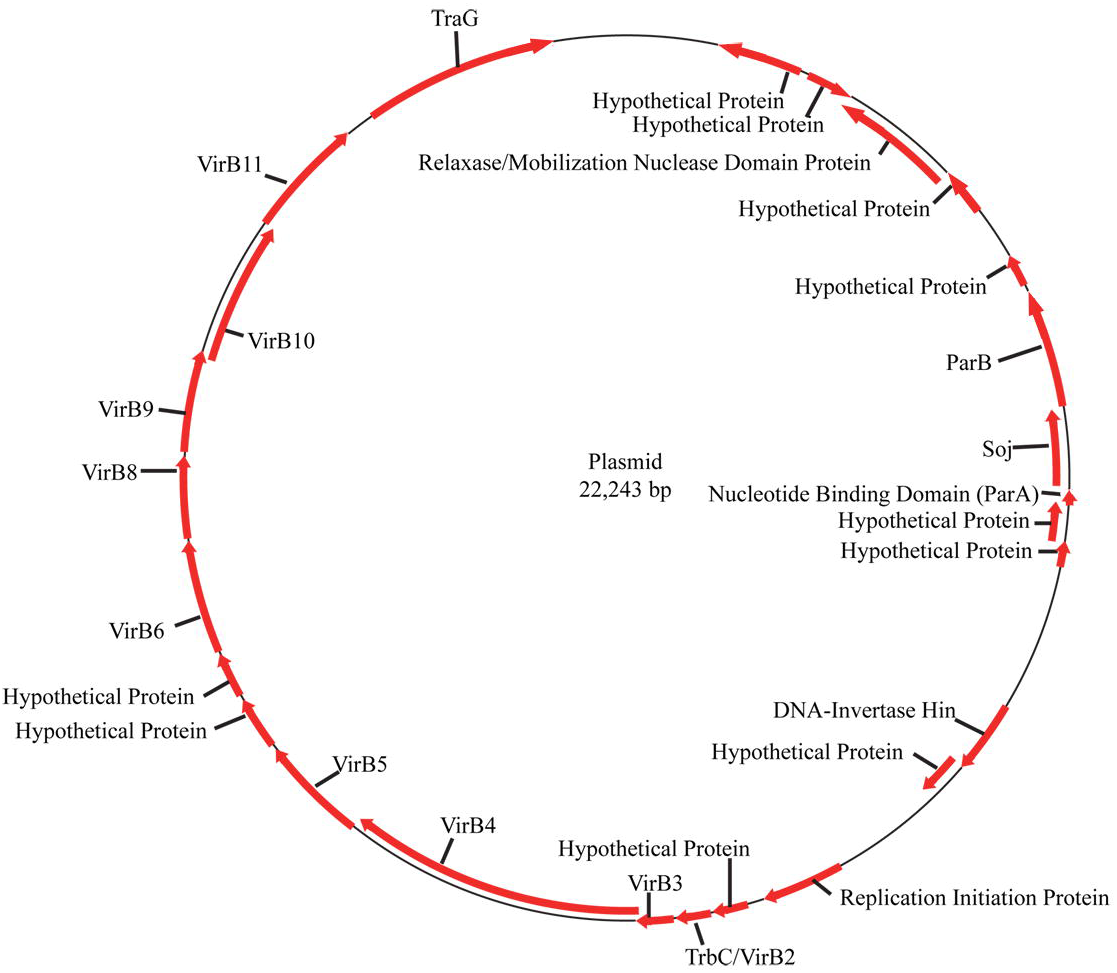

The genomic data showed genes associated with flagellar assembly and function (*fla*G, *flg*A, B, C, E, F, G, H, I, J, K, L, *flh*A, B, F, *fli*D, E, G, H, K, L, M, N, P, Q, S, T, U, W, *mcp*B, *mcp*S, *mot*B, *mot*D, *pct*C, *pom*A, *swr*C, *tar, ycg*R, and an undefined flagellar motor protein). Also identified were the Che genes (*che*A, B, R, V, W, Y, and Z), which can detect chemical conditions in the environment, and interact with the flagellar motor to help the bacteria move to suitable areas within the environment (35). These genes are essential to locate and move to the microaerobic, reduced sulfur rich areas of the sediment this bacterium needs for sulfur oxidation. An aerotaxis gene (similar to *aer*) was also identified, likely allowing the bacteria to locate the microaerobic areas where sulfur oxidation is most efficiently carried out. When associated with a host, reduced sulfur is made accessible to symbionts by the sulfur mining behavior of the clam; however, bacteria in the free-living population must retain key genes for nutrient location and motility (29). Like the environmentally transferred *R. pachyptila* symbiont, the *T.* cf. *gouldi* symbiont has a full complement of flagellar genes, as well as an array of chemotaxis genes (16). Surprisingly, no magnetotaxis genes were identified by our metagenomic analysis, although magnetosome particles were identified in the symbionts of *T*. cf. *gouldi* (29). As magnetosome particles have been identified, associated genes are likely present in the genome, but could not be identified by the pipeline employed here.

A schematic representation of inferred metabolic capabilities of the *T.* cf. *gouldi* symbiont is presented in Figure 2. The symbiont may not be restricted to thiotrophy, and may be able to use alternative metabolic pathways when reduced sulfur is not available. In culturing experiments, the sulfur oxidizing bacterium *Sedimenticola thiotaurini* SIP-G1 is unable to fix carbon in aerobic conditions, where it must instead rely on heterotrophy (36). A previous phylogenetic study (28) placed the *T.* cf. *gouldi* symbiont in a position near *S. thiotaurini* SIP-G1. Based on this phylogenetic placement and the genes identified by this study, the *T.* cf. *gouldi* symbiont may have similar metabolic capabilities; however, without culturing the bacteria in the lab we cannot validate this theory.

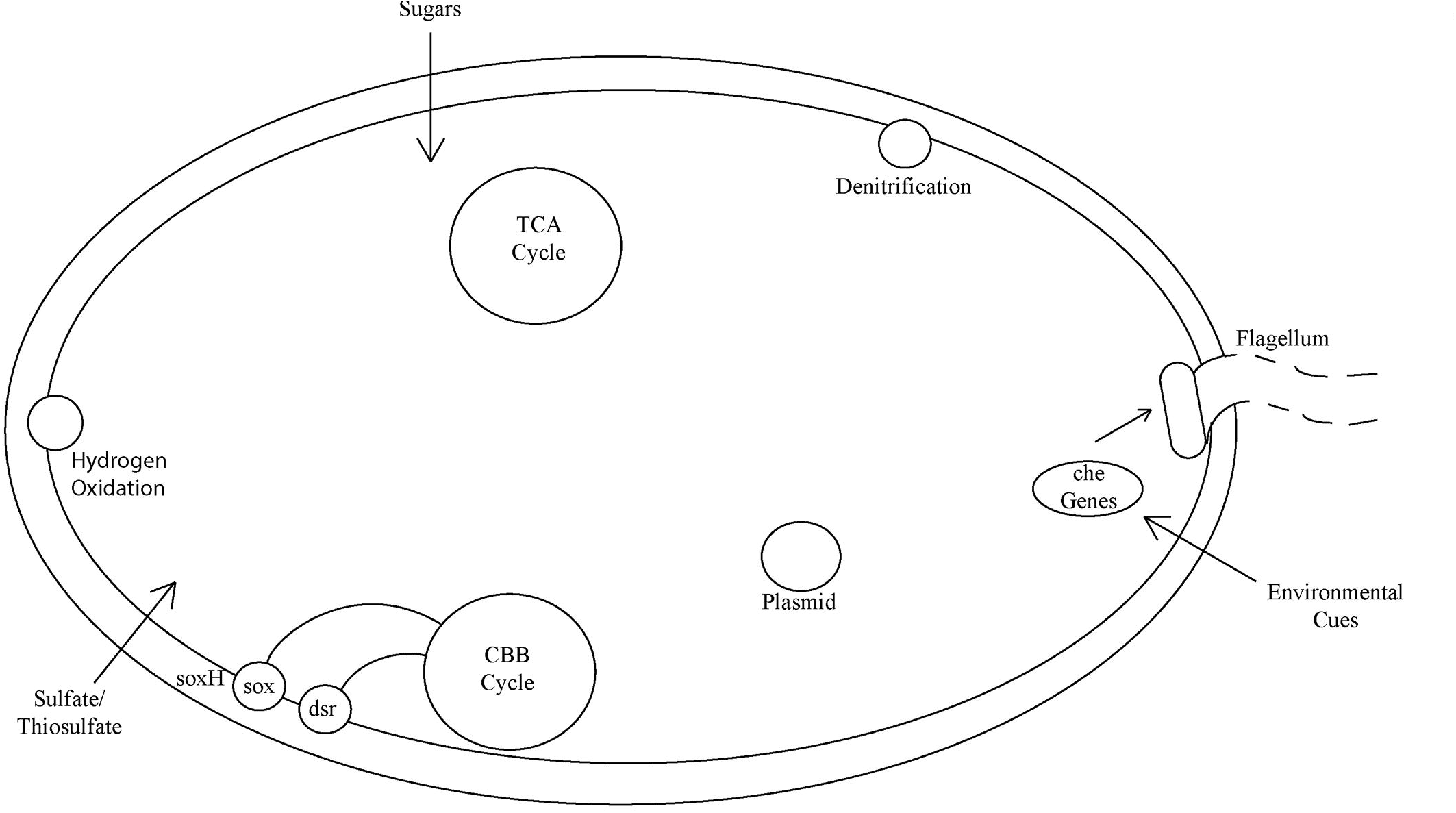

### Amino acid and cofactor synthesis

Symbionts commonly retain genes important for amino acid, vitamin, and cofactor production because the host selects for bacteria that provide the nutrients it requires (15). Many putative gene functions of the *T.* cf. *gouldi* symbiont are involved in amino acid transport and metabolism (229 assignments; Table 1), while cofactor transportation and metabolism are also frequently identified (112 times; Table 1). These functions are also present in free-living bacteria, so while important for the symbiosis, they are also presumably essential to the bacteria outside the host.

### Thioautotrophy

In the *T.* cf. *gouldi* symbiont, the metabolic cycles for carbon fixation and sulfur oxidation are of particular interest. Several genes for the sox and dsr pathways are present (see below), and the symbiont may conduct sulfur oxidation through both these pathways. Both these cycles have been found to function simultaneously in other bivalve chemosymbionts (37, 38). Sulfur compounds within the benthic sediment are patchy, and therefore being able to utilize different forms would increase the habitat range for these bacteria and their bivalve hosts.

*Sox*A, X, Y and Z, which are found in a cluster in the genome of vesicomyid symbionts (37) and form a multi-enzyme system that can oxidize various forms of reduced sulfur (sulfide, thiosulfate, elemental sulfur and sulfite) to sulfate (37, 39) were found in the *T.* cf. *gouldi* metagenome. We found no evidence for soxCD, which is found in some sulfur-oxidizing bacteria but is lacking in others (including in the *Calyptogena* symbiont; 37). The lack of soxCD can manifest itself by the presence of bacterial sulfur globules, which appear as white inclusions in transmission electron micrographs of *T*. cf. *gouldi* symbionts, due to sulfur removal during processing (e.g. Fig. 2B of reference 20). The *T*. cf. *gouldi* symbiont metagenome included *sox*H, a peripheral, thiosulfate inducible sox gene that is located in the periplasm but is not essential for growth on thiosulfate and has an unknown function (40). We also identified *cys*A, shown to import both sulfate and thiosulfate from the environment (41). Adenylylsulfate reductase subunit alpha was found, and its activity was previously detected in thyasirid symbionts (23, 25).

Many of the genes in the dsr cycle were found, with the *dsr*A, B, and C proteins as well as the peripheral *dsr*E suggesting that the pathway is running in an oxidative direction (42). These genes as well as *dsr*K, M, R, S are present in the symbiont genome. An oxidative dsr pathway is present in many well-studied symbionts, including those associated with multiple *Calyptogena* species, *R. pachyptila*, and *Crysomallon squamiferum* (16, 37, 43).

Thirteen putative functions associated with the Calvin-Benson-Bassham Cycle were discovered in the *T.* cf. *gouldi* symbiont (Table 2). The Calvin-Benson-Bassham Cycle in the *T*. cf *gouldi* symbiont utilizes a type II RuBisCO enzyme (20, 28). Chemosymbionts of bivalves often have a reversible pyrophosphate-dependent phosphofructokinase in place of the sedoheptulose-1,7-bisphosphatase that this enzyme replaces, and the fructose 1,6 bisphosphatase genes, which are employed in a reverse TCA cycle (17, 38). However, we were unable to identify any of these three genes in our analysis, but did find ribose 5-phosphate isomerase, which is used in the typical Calvin-Benson-Bassham pathway but is missing in the symbionts of *Calyptogena magnifica* and *R. pachyptila* (15, 16). It is not clear if the thyasirid symbiont has a traditional Calvin-Benson-Bassham cycle, or if the modifications common in other sulfur oxidizing symbionts are also present in this symbiont (17, 38).

### Hydrogen Oxidation

The symbiont also appears capable of hydrogen oxidation using the NAD^+^-reducing hydrogenase *hox*HYUF, and a second set of closely related genes identified as the alpha, beta, delta, and gamma subunits of *hox*S. The enzyme produced by these complexes is bi-directional. It has been described previously in the symbiont of some vestimentiferan worms (44) as well as free-living *Sedimenticola selenatireducens* (45).

### Heterotrophy

Genes associated with the tricarboxylic acid (TCA) cycle were also identified in the *T.* cf. *gouldi* symbiont (Table 3). Interestingly, the TCA cycle in this symbiont does not appear to use the oxoglutarate shunt, and contains both the α ketoglutarate dehydrogenase and citrate synthase enzymes which are commonly lost in chemosymbiotic bacteria and cause the loss of heterotrophic abilities (17). All genes for a functional TCA cycle have been found in the chemosymbiont of *Solemya velum*, which may occur outside of its host (17). The genome of the *R. pachyptila* symbiont also encodes a complete TCA cycle, and contains evidence for response to carbon compounds in the environment, suggesting that it can survive heterotrophically outside the host (16). Sugar phosphotransferase systems (PTS) were also identified in our dataset. These systems can import sugars from the environment, increasing the evidence for some heterotrophic ability. Sugar PTS were identified for fructose, mannose, galactose, and sucrose, suggesting that these substrates can be acquired from the environment, supplementing carbon fixation. In pure culture, the sediment bacterium *S. thiotaurini* SIP-G1 is unable to grow on sulfur oxidation alone, and must be provided with heterotrophic nutrients (36). A similar system may exist within the *T.* cf. *gouldi* symbiont, with heterotrophic growth occurring when environmental conditions are unfavorable for carbon fixation. The ability to utilise multiple carbon sources would be very beneficial during a free-living stage, especially in fluctuating environments where sulfur compounds can be scarce.

**Table 3.** Putative gene functions involved in the TCA cycle.

| Putative gene function | Number of copies | Percent identity | Alignment length (bp) |
| --- | --- | --- | --- |
| 2-oxoglutarate dehydrogenase E1 component | 1 | 70.33 | 209 |
| Aconitate hydratase | 5 | 65.53 | 109 |
| Aconitate hydratase 2 | 1 | 72.11 | 190 |
| Citrate synthase (si) | 1 | 66.97 | 109 |
| Dihydrolipoamide dehydrogenase | 2 | 73.13 | 89 |
| Dihydrolipoamide dehydrogenase of pyruvate dehydrogenase complex | 1 | 75.83 | 211 |
| Dihydrolipoamide succinyltransferase component (E2) of 2-oxoglutarate dehydrogenase complex | 2 | 75 | 138.5 |
| Fumarate hydratase class I, aerobic | 2 | 88.05 | 163 |
| Isocitrate dehydrogenase [NADP] | 3 | 69.21 | 265.33 |
| Malate dehydrogenase | 2 | 63.59 | 209 |
| Succinate dehydrogenase flavoprotein subunit | 4 | 77.19 | 137.5 |
| Succinate dehydrogenase iron-sulfur protein | 1 | 70.83 | 144 |
| Succinyl-CoA ligase [ADP-forming] alpha chain | 3 | 81.71 | 185 |
| Succinyl-CoA ligase [ADP-forming] beta chain | 4 | 72.13 | 178 |

### Anaerobic respiration

The *T.* cf. *gouldi* symbiont appears to be capable of performing denitrification, as the genes for the nar and nos pathways are present in the genome. Denitrification is the process that reduces potentially harmful nitrogen compounds (nitrates, nitrites, and nitric oxide) into harmless, inert N2 through anaerobic respiration. Denitrification may provide multiple advantages to both host and symbiont, in addition to allowing bacterial ATP synthesis. First, by reducing harmful nitrogenous compounds, the bacteria protect both themselves and their host from toxic effects. Second, by decreasing the symbiont’s oxygen requirements, there is less competition with the host for this limited resource in the thyasirid’s endobenthic environment. Third, the pathways could allow the bacteria to respire anaerobically in anoxic sediments, and therefore broaden the organism’s free-living niche. Notably, the closely related free-living bacterium *S. thiotaurini* SIP-G1 from salt marsh sediments is capable of anaerobic respiration using nitrate and nitrite, but can also grow under hypoxic conditions (36). Dissimilatory nitrate respiration genes have also been identified in the symbionts of *Vesicomyosocius okutanii, R. pachyptila*, and *Bathymodiolus thermophilus* (16, 46, 47).

Recent work has shown some sulfur-oxidizing chemosymbionts can also fix atmospheric nitrogen into bioavailable forms (18, 48), however, we did not find any evidence of N fixation genes in the *T.* cf. *gouldi* symbiont. The closely-related free-living bacterium *S. thiotaurini* SIP-G1 does however have a complete nitrogen fixation pathway (36).

## Conclusions

The genomic data collected on the *T.* cf. *gouldi* symbiont corroborates the previous data suggesting a facultative relationship, with the host clams being inoculated from the environment (28, 29). The timing of this inoculation during the host’s lifespan is still unclear and further research is needed to determine when the host is competent for symbiont uptake. The symbiont population is a collection of closely related individuals, although the population is not clonal and some variation is present. There is no evidence of genome reduction in these symbionts, and the genomic data supports evidence of an environmental (non-symbiotic) habitat. In particular, the presence of a functional flagellum and chemosensory abilities supports the presence of a free-living population, as reported previously (29).

The metabolic capabilities of the symbionts are comparable to previously described sulfur oxidizing bacteria. The symbionts utilize multiple pathways for sulfur oxidation, both sox and dsr, and the Calvin-Benson-Bassham Cycle for carbon fixation. The denitrification pathway that is also present would allow for carbon fixation in anaerobic areas; when outside the host, sulfides are predominantly found in micro-oxic areas. Unlike many obligate symbionts, the thyasirid symbiont appears to have a functional TCA cycle and sugar importers allowing it to be heterotrophic. The bacteria may utilize autotrophy or heterotrophy under different conditions, like *S. thiotaurini* SIP-G1 (36).

Further research into the thyasirid symbiont genome may be beneficial in tracking the changes required for life as a bivalve symbiont, and experimental studies could reveal whether symbionts are capable of reverting to a non-symbiotic state after they have become associated with their host. The *T.* cf. *gouldi* symbiosis provides a unique opportunity to investigate how symbioses evolve as this appears to be a relatively less derived and interdependent relationship compared to other bivalve symbioses which are intracellular. More research into the metabolic capabilities of the symbiont and how they interact with the host would provide insights into how this relationship has evolved, and the mechanisms that allow it to be maintained. Comparing the different symbiont phylotypes capable of associating with a single host species would also improve our understanding of this relationship and of the potential benefits of flexible host-symbiont pairings.

## Methods

### Sample Collection and Sequencing

Sediment was collected in August 2010 using a Petersen grab from Neddy’s Harbour, in the fjord of Bonne Bay, Newfoundland, Canada (49°31’21.44"N, 57°52’11.07"W), at a depth of roughly 15 m. Sediment was wet sieved using a 1 mm mesh and specimens of *T.* cf. *gouldi* were collected and transported to Memorial University, St. John’s, Newfoundland. Total DNA was extracted from the gills of a single individual (host OTU 1; reference 20) using a Qiagen Blood and Tissue Kit and stored at −20°C in the elution buffer provided. Before sequencing, total DNA was transferred to nuclease free water. An Ion Torrent Fragmentation Kit was used and fragments of approximately 200 bp were selected using gel size selection and extraction (Qiagen Gel Extraction Kit), purified (Qiagen DNA Purification Kit) following manufacturer’s instructions, and concentrations assessed with an Agilent Bioanalyser. Sequencing was conducted on an Ion Torrent PGM Sequencer following the manufacturer’s protocols (V2.2). A 316 chip was used for sequencing. Due to poor load rates, two sequencing runs were conducted and the data were combined before further processing.

### Assembly and Annotation

Reads were quality checked and trimmed using FastQC, FAstQ Groomer, FastQ Quality Trimmer, and Filter FastQ the Galaxy Website (usegalaxy.org) and FastQC software (49, 50); any reads less than 50 bp long were removed at this stage. A quality score of 20 was used, allowing one base below the cutoff score within the read, and trimming was conducted on both ends. Filtered reads were binned using MEGAN5 (51).

Assembly of the binned data was conducted using SPAdes (52). Ion Torrent specific settings with kmers 27, 35, 55 and 77 were used. All bacterial, unknown, and unassigned reads were used for assembly. Contigs of 200 bp or more were entered into the pipeline. Annotation was run using the MG-RAST website (http://metagenomics.anl.gov/) (53), the RefSeq, KOG, and Subsystems databases were used with the e-value cut-off set at 5, % identity 60, min length 15, and min abundance 1. A secondary annotation was conducted using PROKKA (54).

## Acknowledgements

This research was funded by Natural Sciences and Engineering Research Council (NSERC) Discovery Grants (RGPIN 386087-2010 and 06548-2015) to SCD, and an NSERC Postgraduate Scholarship – Doctoral to BM.

We would also like to thank the Lang lab, specifically Yunyun Fu, for training and assistance with the Ion Torrent PGM.

